# Phosphorylated paxillin and FAK constitute subregions within focal adhesions

**DOI:** 10.1101/2020.09.24.311126

**Authors:** Michael Bachmann, Artiom Skripka, Bernhard Wehrle-Haller, Martin Bastmeyer

## Abstract

Integrin-mediated adhesions are convergence points of multiple signaling pathways. Their inner structure and their diverse functions can be studied with super-resolution microscopy. We used structured illumination microscopy (SIM) to analyze spatial organization of paxillin phosphorylation (pPax) within adhesions. We found that pPax and focal adhesion kinase (FAK) form spot-like, spatially defined clusters within adhesions in several cell lines. In contrast, other adhesion proteins showed no consistent organization in such clusters. Live-cell super-resolution imaging revealed that pPax-FAK clusters persist over time but modify distance to each other dynamically. Moreover, we show that the distance between separate clusters of pPax is mechanosensitive. Thus, in this work we introduce a new structural organization within focal adhesions and demonstrate its regulation and dynamics.

## Introduction

Integrin-mediated adhesions between cells and the surrounding extracellular matrix are not only important for physical anchorage but are also converging points for different intra- and extracellular signals (Bachmann, Kukkurainen, Hytönen, & Wehrle-Haller, 2019; Conway & Jacquemet, 2019; Green & Brown, 2019). These adhesions consist of a plethora of proteins establishing the so-called adhesome (Byron, Humphries, Bass, Knight, & Humphries, 2011; Kuo, Han, Hsiao, Yates, & Waterman, 2011; Schiller, Friedel, Boulegue, & Fässler, 2011). Based on these adhesome studies, a meta-analysis curated a consensus adhesome whose members form stable clusters connected with each other into a complex network (Horton et al., 2015). Advent of super-resolution microscopy confirmed these clusters by showing that members of a cluster are organized within the same axial layer (Kanchanawong et al., 2010). While axial layering is now comparably well documented, it remains less explored if or how a lateral structure is implemented within adhesions. So far, pointillistic super-resolution methods revealed the existence of sub-clusters within adhesions (Bachmann, Fiederling, & Bastmeyer, 2016; Changede, Xu, Margadant, & Sheetz, 2015; Shroff, Galbraith, Galbraith, & Betzig, 2008; Shroff et al., 2007), but a connection between spatial organization and function (as in the case for axial organization) remains to be shown.

One of the most important proteins for structure and function of integrin-mediated adhesions is paxillin. Paxillin, part of the LIM-domain protein family, is recruited to adhesions by the integrin-activators talin and kindlin (Pinon et al., 2014; Theodosiou et al., 2016), facilitated by membrane interaction of LIM4 domain (Ripamonti, Liaudet, & Wehrle-Haller, 2020). Integrin activation and integrin-mediated cell adhesion can be realized in the absence of paxillin but cell spreading and proliferation relies on paxillin recruitment to integrin adhesions (Pinon et al., 2014; Soto-Ribeiro et al., 2019). Structurally, paxillin serves as a scaffolding protein whose multiple binding partners can be regulated by paxillin phosphorylation (Deakin & Turner, 2008). Tyrosine-phosphorylation at Y31 (pPax-Y31; one letter amino acid code: Y for tyrosine) and Y118 (pPax-Y118) by a complex of focal adhesions kinase (FAK) and Src has been intensively studied and appears to recruit activators and inhibitors for Rho-GTPases (Deakin & Turner, 2008; Petit et al., 2000; Schaller & Parsons, 1995; Thomas et al., 1999). These phospho-tyrosine sites are also important for adhesion dynamics as demonstrated with phosphomimetic and none-phosphorylatable mutants for Y31 and Y118 (Ripamonti et al., 2020; Zaidel-Bar, Milo, Kam, & Geiger, 2007). Interestingly, reduced dynamics as observed with none-phosphorylatable paxillin is also observed with wt-paxillin when FAK is kept inactive or when FAK is genetically depleted (Ilić et al., 1995; Swaminathan, Fischer, & Waterman, 2016; Webb et al., 2004). These data highlight the dependence of pPax-Y31/Y118 signaling on FAK presence and activation status. Fittingly, adhesome studies mentioned above cluster FAK and paxillin within the same group (Horton et al., 2015). Moreover, paxillin and FAK localize in the same axial layer as demonstrated by super-resolution microscopy (Kanchanawong et al., 2010), and form a complex preferentially when paxillin is phosphorylated at Y31 and Y118 (Choi, Zareno, Digman, Gratton, & Horwitz, 2011; Digman, Brown, Horwitz, Mantulin, & Gratton, 2008).

Here, we studied the lateral organization within integrin-mediated adhesions with super-resolution structured illumination microscopy (SIM). We analyzed several adhesome proteins belonging to different clusters or axial layers, respectively. We found different spatial organization of adhesome proteins within the same adhesion. Most proteins, including paxillin, were distributed quite homogenously throughout an adhesion; however, phosphorylated paxillin and FAK showed confinement in distinct clusters. Single adhesions could contain several of these spot-like clusters that were well separated from each other. Moreover, we detected these spots in different cell lines and showed with super-resolution live cell imaging that the spacing of spots to each other is very dynamic. Finally, we demonstrated that this spacing is force-sensitive and dependent on vinculin recruitment. These findings demonstrate that integrin-mediated adhesions show a lateral structuring for pPax and FAK under a variety of conditions that can be regulated in an active and force-dependent manner.

## Results

### Substructure of adhesome proteins in focal adhesions

Rat embryonic fibroblast (REF) cells are an established cell line for the analysis of adhesions and their composition (Cavalcanti-Adam et al., 2007; Franz & Müller, 2005; Gudzenko & Franz, 2015; Hoffmann, Fermin, Stricker, Ickstadt, & Zamir, 2014). We cultured these cells on fibronectin-coated cover slips and transfected fluorescently tagged proteins of interest or performed secondary immunostaining to analyze several consensus adhesome proteins. We tested (i) two different integrins (β3-GFP and BMB5 antibody against α5, staining α5β1 or αVβ3 respectively), (ii) the integrin activators talin-1 and kindlin-2, (iii) paxillin, FAK, and pPax-Y118 as indicators for adhesion mediated signaling, and (iv) vinculin and zyxin as adhesome components involved in adhesion-cytoskeleton linkage, as well as (v) actin itself. Imaging was performed with SIM, allowing us to analyze the organization of focal adhesions with two times higher resolution compared to diffraction-limited microscopic methods (Bachmann et al., 2016). To analyze the spatial organization of adhesome proteins inside single cell-matrix adhesions we focused on focal adhesions (Gardel, Schneider, Aratyn-Schaus, & Waterman, 2010). With the experimental conditions we used, we rarely observed fibrillar adhesions, while nascent adhesions or focal complexes are too small to manifest a sub-structural organization that could be detected with SIM. Due to SIM, we could confirm the earlier observation by different groups that focal adhesions are split into parallel ‘stripes’ (as indicated for example by the paxillin staining in Fig. 1), compared to a more homogenous organization observed with diffraction-limited microscopy (Hu et al., 2015; Young & Higgs, 2018). We found that these stripes were well separated from each other, and could be considered as individual focal adhesions in our experimental setup. A closer analysis of protein localization in these focal adhesions revealed that some adhesome proteins were organized in a distinct substructure. Intensity profiles along the long axis of focal adhesions showed that especially pPax-Y118 and FAK were segregated into separate clusters within the same adhesion. Especially pPax-Y118 spots showed a remarkable regularity concerning the spacing of these clusters. In contrast, paxillin, vinculin, zyxin, and actin staining intensities varied along the focal adhesions, but a continuous staining was present throughout their whole length; separated spatial clusters were only visible occasionally and not as regularly as observed for pPax-Y118 and FAK. Fluorescent stainings of the remaining proteins – α5β1 and αVβ3 integrin, talin-1, and kindlin-2 – revealed a behavior somewhat between the two cases mentioned above. This is highlighted by the zoom-in for talin-1 in Figure 1 which shows focal adhesions with continuous and spot-like organization of talin-1 next to each other.

**Figure 1:**
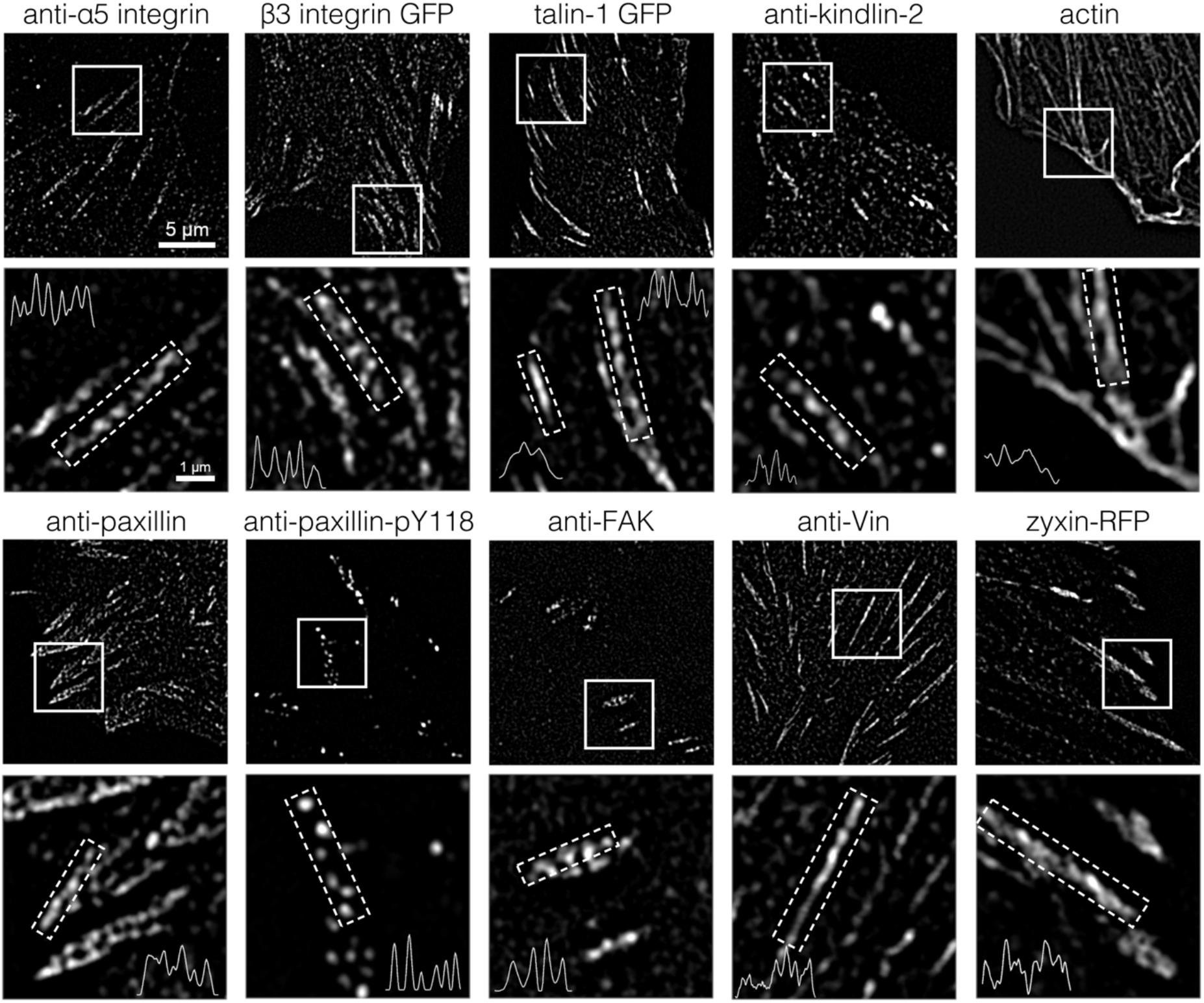
Analysis of spatial organization of adhesome proteins within focal adhesions. Rat embryonic fibroblast (REF) cells were cultured on fibronectin-coated cover slips. Cells were either transfected for the indicated protein or stained with indirect immunofluorescence or with phalloidin for actin. Imaging was performed with SIM. White boxes indicate zoom-in regions shown below. The intensity profiles shown in the zoom-ins are calculated for adhesions within boxes with dotted white lines. Please note that variations in intensity along the intensity profiles are observed for all proteins. However, some conditions revealed distinct intensity spots indicating well separated protein clusters; pPax-Y118 and FAK show this organization most consistently. All overview images and zoom-ins are shown with the same magnification as indicated by the scale bar.

Thus, SIM revealed a spatial organization within focal adhesions that differed not only between different adhesome proteins but even relied on phosphorylation status, as shown for pPax-Y118 compared to paxillin.

### Phosphorylated paxillin is spatially restricted to separated clusters

Surprised by the difference in paxillin vs. pPax-Y118 organization we decided to analyze their spatial organization in detail and in a quantitative manner. We cultured REF cells on cover slips and stained endogenous paxillin as well as pPax-Y118 (Fig. 2A). Magnifications of single focal adhesions confirmed our earlier observation that paxillin is spread rather homogenously throughout adhesions, while pPax-Y118 shows a spatial organization in discrete clusters (Fig. 2B-D). To analyze the pPax-Y118 spacing in more detail, we developed a custom-written Matlab code that measures the center-to-center distance of protein clusters based on the position of their intensity maxima (Fig. S1). The analysis revealed a distribution of pPax-Y118 cluster distances with an average separation around 508 nm (Fig. 2E, G). The algorithm also allowed to extract an average cluster diameter of 210 +/− 70 nm (Fig. 2F).

**Figure 2:**
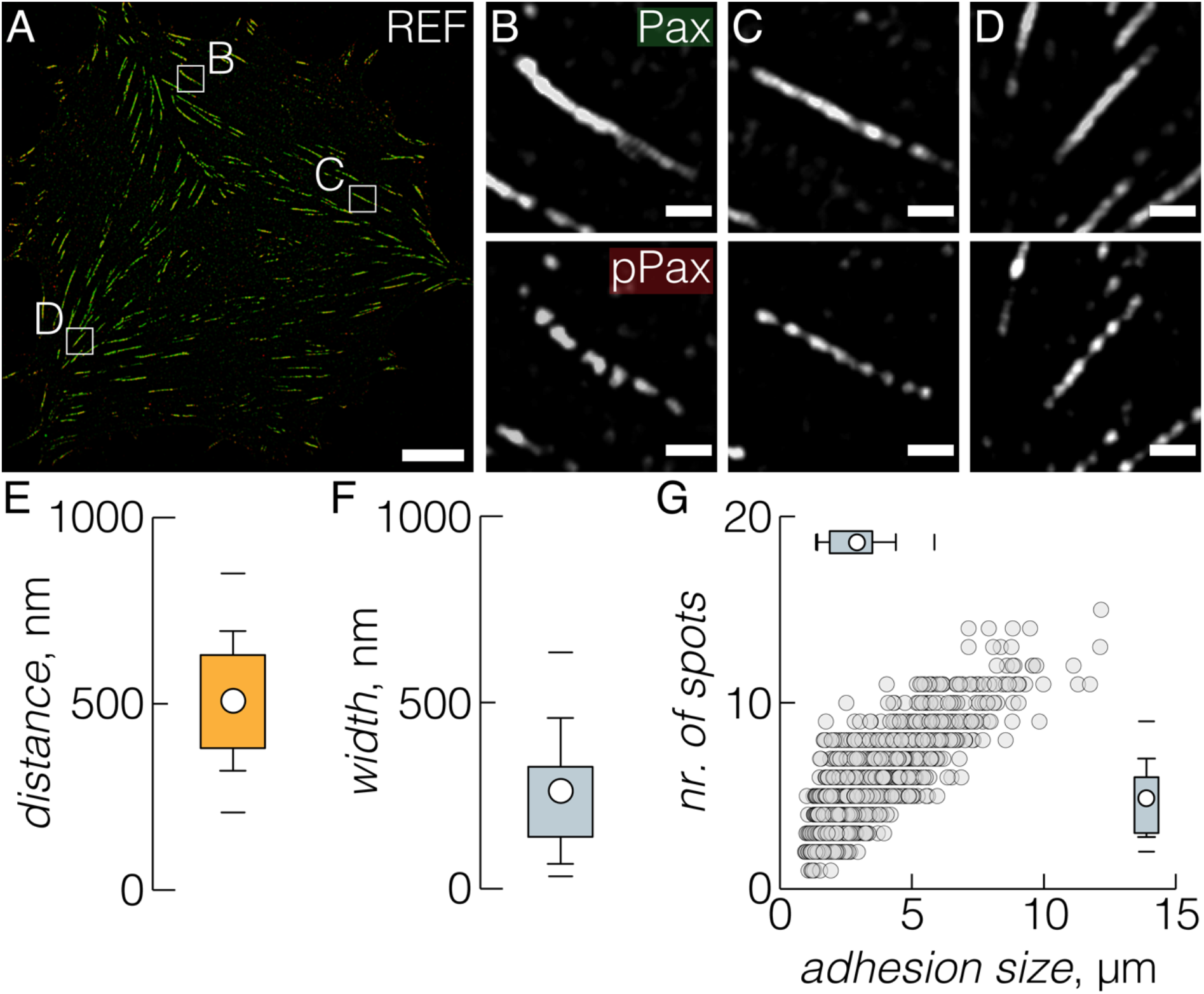
Analysis of inner organization of focal adhesions for paxillin and pPax-Y118. (A) REF cell was cultured on fibronectin-coated cover slip and stained for paxillin (green) and pPax-Y118 (red). (B-D) Zoom-ins for paxillin staining or pPax staining indicate that pPax-Y118, but not paxillin, is organized in discrete spatial clusters. (E) Analysis of pPax-Y118 cluster distances and (F) cluster width based on a custom-written Matlab code (see material & method section and Fig. S1; N = 3, n = 15; circle in box plots indicates mean, see material & method section for detailed explanation). (G) Distribution of number of clusters within a single adhesion vs. the length of the respective adhesion. Every dot represents one adhesion. Box plots at the side indicate distribution of number of clusters or adhesion size, respectively. Scale bars: 10 μm in overview, 1 μm in zoom-ins.

While the size of these clusters is close to the resolution limit of most microscopes, spacing between spots should be well resolved by diffraction-limited microscopes. Therefore, we imaged paxillin and pPax-Y118 in the same focal adhesions with confocal laser scanning microscopy (LSM), total internal reflection microscopy (TIRF), and SIM (Fig. S2A, B). Direct comparison revealed that only SIM consistently resolved separated pPax-Y118 clusters. Meanwhile, LSM and TIRF were able to resolve some of the pPax-Y118 clusters that were observed with SIM. Thus, diffraction-limited microscopy indicates pPax-Y118 clusters to a certain degree confirming that this spatial pPax-Y118 organization is not a microscopic artifact. Additionally, we imaged pPax-Y118 stained cells with AiryScan super-resolution mode (1.7x resolution improvement compared to 2x of SIM, (Jacquemet, Carisey, Hamidi, Henriques, & Leterrier, 2020)). AiryScan, as SIM, was able to resolve an organization of pPax-Y118 in spots (Fig. S2C). Thus, resolution improvement of at least 1.7x consistently indicates clusters of pPax-Y118 within single adhesions independent of the specific microscopic technique used.

Furthermore, to confirm that differences in spatial organization are not induced by staining pitfalls, like unspecific antibody binding, we performed titration experiments of primary antibodies for paxillin (Fig. S4) and pPax-Y118 (Fig. S3). We could not detect a concentration dependent effect of antibodies on the appearance of paxillin compared to pPax-Y118. Importantly, we also tested a primary antibody for pPax-Y118 from another supplier and an antibody for pPax-Y31 (Fig. S3). Both antibodies against pPax-Y118 confirmed the observation that pPax-Y118 organizes in discrete clusters, as does pPax-Y31.

### Phosphorylated tyrosines in focal adhesions organize in clusters

Besides pPax (Y31 and Y118), we observed that FAK also organizes in discrete clusters with a near-regular spacing (Fig. 1). Interestingly, FAK was reported to complex preferentially with paxillin phosphorylated at Y31 and Y118 (Choi et al., 2011; Digman et al., 2008). In order to phosphorylate paxillin Y31 and Y118, FAK has to be autophosphorylated at its tyrosine 397 (pFAK) which is a marker of activated FAK. Thus, we wondered about the spatial organization of pFAK within focal adhesions. To test for this, we performed double stainings of focal adhesions in REF cells for paxillin and pFAK (Fig. 3B) as well as paxillin and pPax-Y118 as control (Fig. 3A). SIM images revealed that pFAK (Fig. 3B) is spatially organized in clusters as seen before for FAK (Fig. 1) and for pPax-Y118 (Fig. 2 and 3A). We went on and stained with primary antibodies that bind promiscuously to phosphorylated tyrosines (pTyr; Fig. 3C). Interestingly, this staining also revealed a spatial pattern of separated clusters. Furthermore, a quantification of the distances between pFAK and pTyr clusters matched closely the average spacing observed for pPax-Y31 and pPax-Y118 (Fig. 3D; pFAK: 508 nm, pTyr: 507 nm; pPax: around 510 nm in all cases, see Fig. 2 and Fig. S3). Thus, spot-like substructures of clusters within focal adhesions emerge as a common feature of the signaling proteins pPax and pFAK, and might be generalized to focal adhesion proteins with phosphorylated tyrosines.

**Figure 3:**
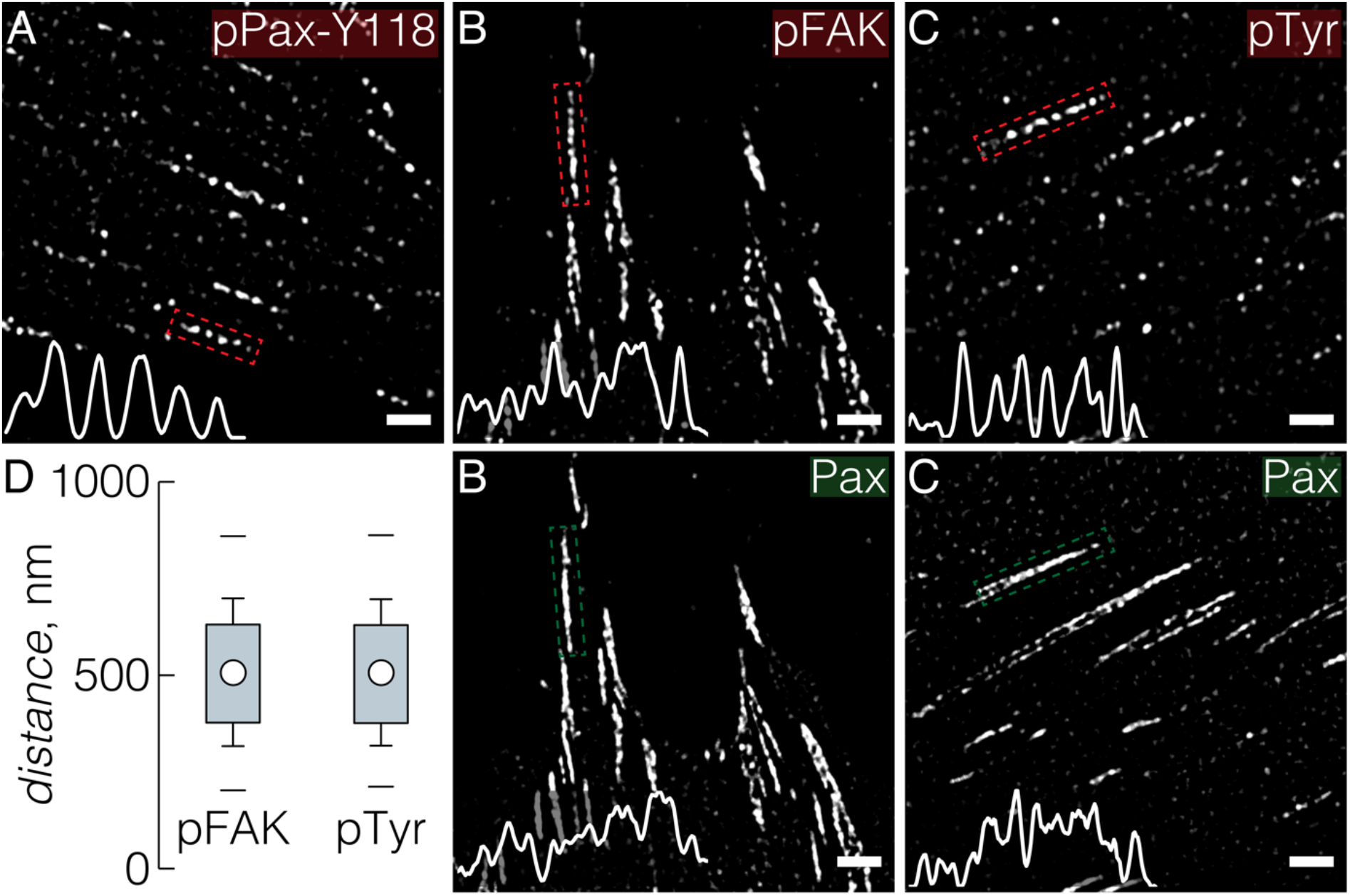
Organization in separated clusters is observed for adhesome proteins with phosphorylated tyrosines. (A) Zoom-in with intensity profile for REF cell stained for pPax-Y118 as described in Fig. 2. Please note the regularly spaced pPax-Y118 clusters (Line profile from adhesion in dotted red box). (B) REF cell treated as in (A) was stained for paxillin (green) and pFAK (red). Staining and intensity profiles indicate organization of pFAK in separate clusters compared to paxillin. (C) REF cell treated as in (A) was stained for paxillin (green) and pTyr (red) again indicating cluster like organization of proteins with phosphorylated tyrosines. (D) Distance analysis of pFAK (N = 3, n = 38) and pTyr (N = 3, n = 38) clusters shows a similar distribution as pPax-Y118 (see Fig. 2E). Scale bars: 2 μm.

### pPax organizes in spatially separated clusters in several cell lines

Next, we tested whether the observation of pPax-Y118 organization in separated clusters can be extended to different cell lines. We compared other established fibroblast cell lines (mouse embryonic fibroblasts, MEFs; mouse fibroblasts, NIH 3T3; primary human foreskin fibroblasts, HFF), a cancer cell line (HeLa), and an epithelial cell line (normal rat kidney cells, NRK) (Fig. S5). In all cell lines, pPax-Y118 showed an organization in separated clusters in contrast to paxillin (Fig. S5A’-E’). Quantitative analysis revealed that pPax-Y118 cluster spacing is overall conserved between different cell lines (Fig. S5 F,G). In sum, all experiments so far confirmed that segregated clusters of FAK, of phosphorylated tyrosines in adhesion proteins, and especially of pPax-Y118, are a consistent feature of lateral organization within single focal adhesions.

### Spatial organization of pPax and FAK is dynamic while remaining in clusters

To understand the spatial organization of pPax and FAK in more detail we set out to analyze the temporal evolution of this pattern in focal adhesions. First, we cultured REF cells on cover slips and fixed them at different time points after cell seeding, followed by immunostainings for paxillin and pPax-Y118 (Fig. 4 A,B). Starting with culturing cells for 2 hrs, we observed focal adhesions with the same spatial organization as described above: paxillin is spread throughout adhesions while pPax-Y118 is organized in distinct spatial clusters. Quantitative analysis revealed that the spacing between neighbored pPax-Y118 clusters is preserved over time and always remains around 501 nm (Fig. 4C). This experiment, however, measured pPax-Y118 spacing as an average for many focal adhesions from different cells. Thus, the possibility remained that single clusters of pPax are dynamic and change their spacing over time. Live-cell SIM would be ideal to analyze the origin and development of pPax clusters in more detail. However, live-cell imaging of protein phosphorylation is challenging. Instead, we made advantage of the fact that FAK also showed an organization in discrete clusters (Fig. 1). We transfected REF cells with FAK-GFP and the actin marker F-tractin tdTomato, and monitored living cells with SIM at a 1 frame/min rate (Fig. 5A, Movie 1). We observed that FAK is organized in distinct spatial clusters directly after adhesion initiation at a filopodia – matrix interface (compare time point 0 and 1 min in Fig. 5A” and 5A”’, Movie 2). Following FAK-GFP (Fig. 5A”’) also showed that FAK remains in separate clusters most of the time. However, it was not possible to follow a single spot of FAK and its development between two time points indicating insufficient temporal resolution. Therefore, we monitored only FAK-GFP in REF cells allowing us to increase the temporal resolution to 1 frame every 15 seconds (Fig. 5B, Movie 3 and 4). With this frame rate we observed merging and splitting of two neighboring FAK-GFP clusters while new clusters emerged or disappeared. But even at this temporal resolution it remained challenging to follow the development of individual clusters of FAK-GFP, pointing to a very dynamic regulation of the spatial organization of FAK.

**Figure 4:**
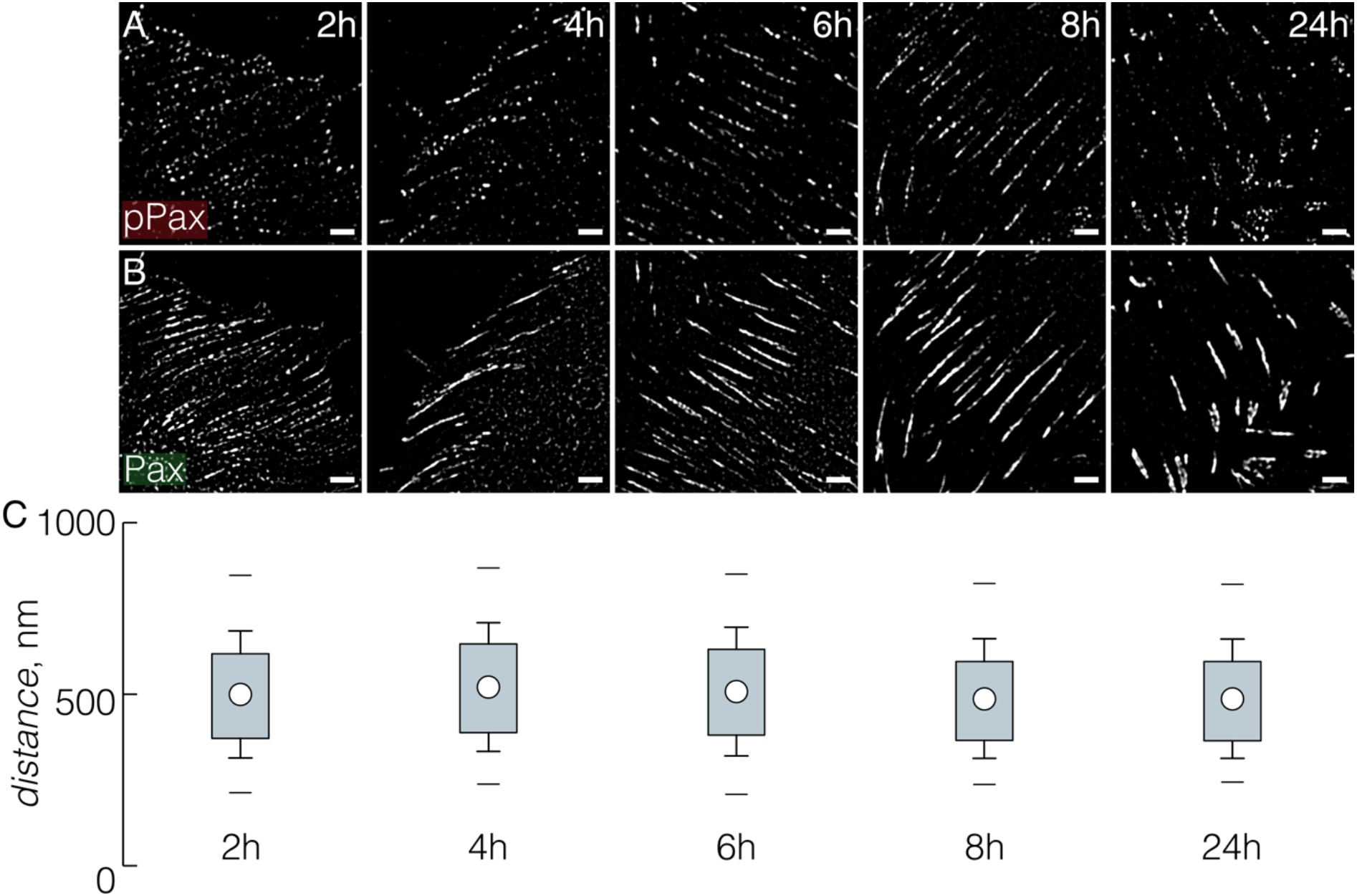
Average distances between neighbored pPax-Y118 clusters show no time dependency. (A) REF cells were cultured on fibronectin-coated coverslips and fixed at the indicated time point and stained for paxillin (green) and pPax-Y118 (red). (A) pPax-Y118 staining revealed spot-like patterns for all time points while (B) paxillin staining persisted throughout the long axis of focal adhesions. (C) Distance analysis for all time points showed no significant differences in pPax-Y118 spacing for time points analyzed (N=3, at least 15 cells per condition). No significant changes were detected (see material & method section). Scale bar: 2 μm.

**Figure 5:**
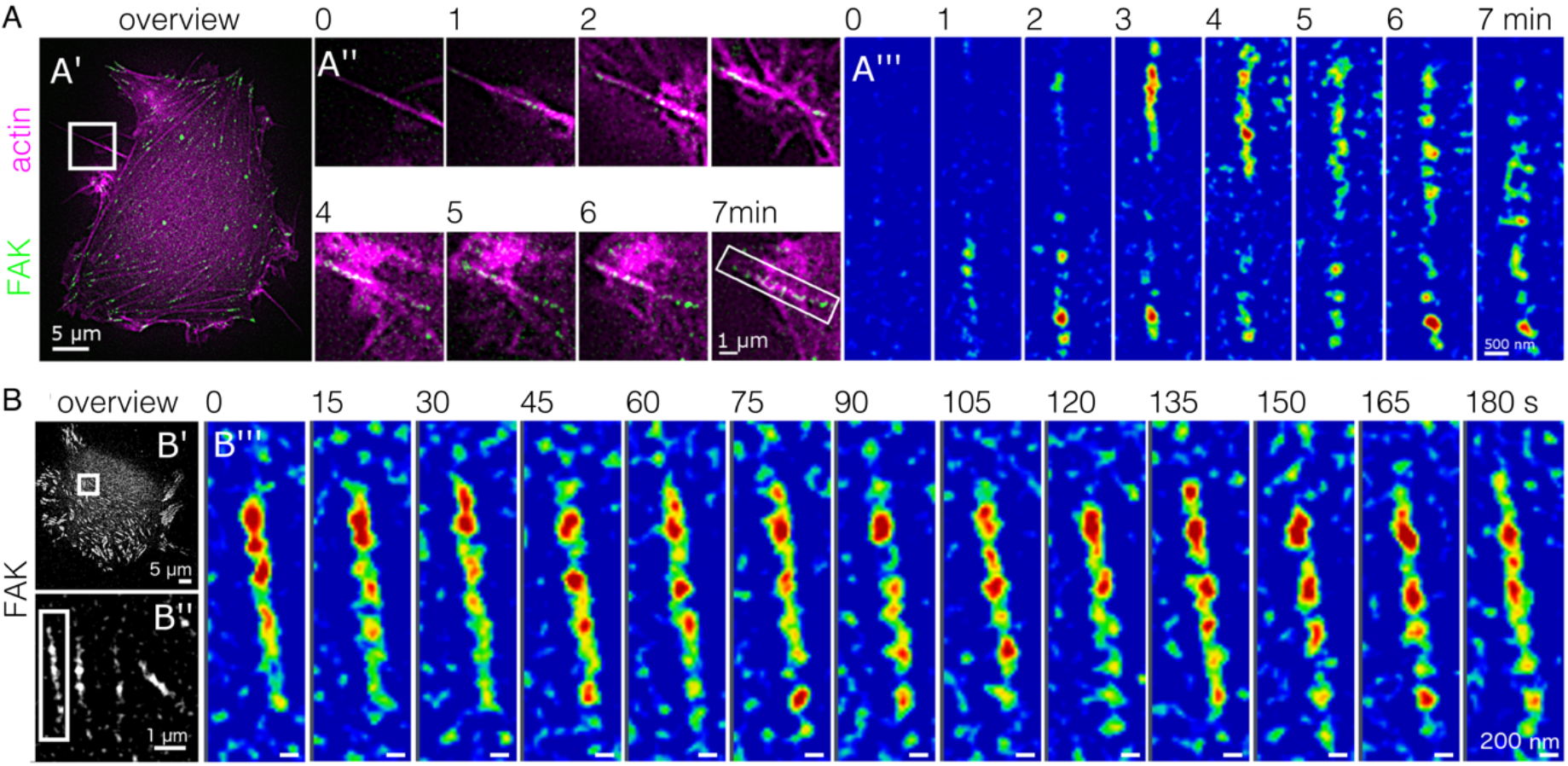
Clusters of FAK persist over time but rearrange dynamically. (A) REF cells were transfected for FAK-GFP and the actin marker F-Tractin tdTomato. Living cells were monitored with SIM with a frame rate of one SIM image per minute. (A’) Magnifications of the white box in the overview image show the development of nascent adhesions to mature focal adhesions (see also Movie 1). (A”) Overlay images of FAK (green) and actin (magenta) and (A”’) false-color coded images of FAK (see also Movie 2). Please note that within one minute (time point 0 min to 1 min) FAK directly appears in spatially separated clusters. FAK remains constrained in clusters throughout the analysis. (B-B”) REF cells only transfected with FAK-GFP imaged at one SIM image per 15 seconds (see also Movie 3). (B”’) Zoom-in from B” shows the temporal evolution of FAK-GFP (shown in false-color) in a single focal adhesion (see also Movie 4). FAK-GFP remained in clusters over time while single FAK-spots rearrange dynamically. Scale bars as indicated.

Overall, super-resolution live-cell imaging revealed that organization in clusters occurs early on for FAK during adhesion formation and remains stable over time. However, while this existence of clusters is stable, single FAK clusters in focal adhesions show a very dynamic behavior, exceeding our technical possibilities to fully resolve the development of them.

### Spacing of pPax clusters is mechanosensitive and dependent on vinculin

Finally, we wondered whether it is possible to interfere with the spacing between neighbored pPax clusters. Publications indicated that actomyosin contractility influences paxillin phosphorylation (Pasapera, Schneider, Rericha, Schlaepfer, & Waterman, 2010; Zaidel-Bar et al., 2007). Thus, we treated REF cells with the myosin inhibitor blebbistatin or with Y-27632, an inhibitor of the Rho-ROCK pathway (Fig. 6). Both treatments caused an increase in the number of peripheral nascent adhesions. However, we could still observe focal adhesions and use them for quantitative analysis of pPax-Y118 spacing (Fig. 6A’-C’). Both treatments caused a reduction in pPax-Y118 cluster spacing. Thus, reduced contractility caused a higher density of pPax clusters within focal adhesions.

**Figure 6:**
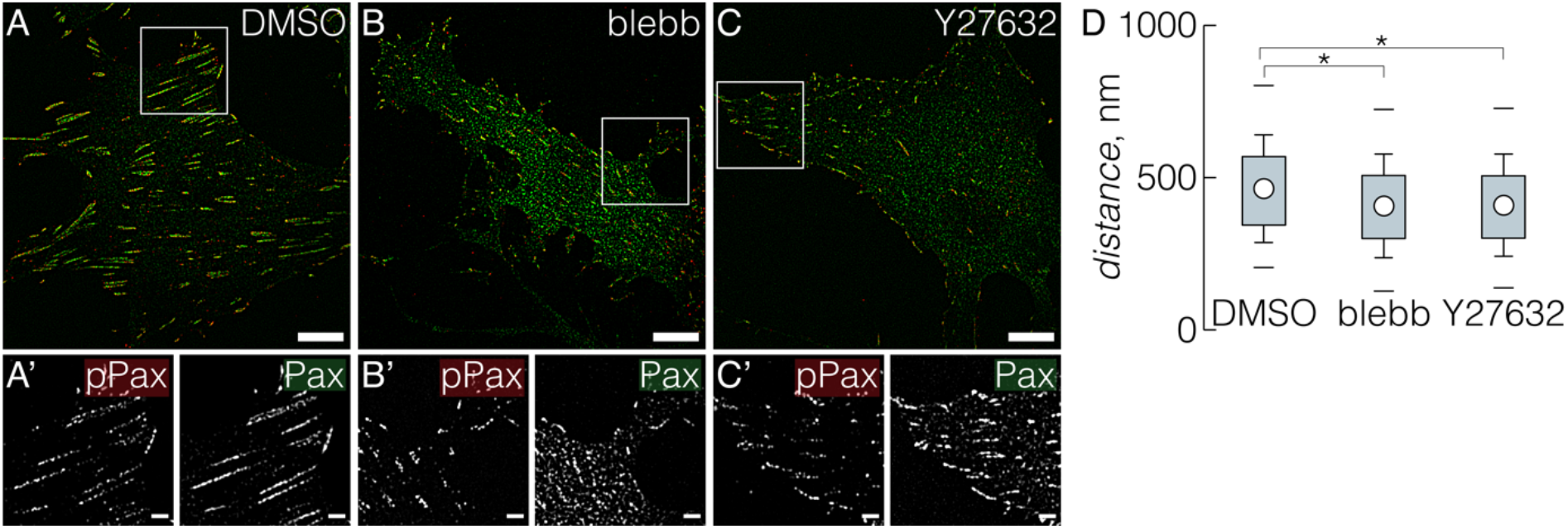
pPax-Y118 cluster distance is mechanosensitive. (A-C) REF cells were cultured for 6 hrs and (when indicated) incubated with 20 μm blebbistatin or Y27632 for the last hour of the experiment. After fixation, cells were fixed and stained for paxillin (green) and pPax-Y118 (red). (A’-C’) Zoom-ins according to the white boxes in (A-C) show pPax-Y118 or paxillin as indicated. (D) Quantitative analysis of pPax-Y118 cluster distance indicates consistent reduction in pPax-Y118 cluster distance in cells with reduced contractility due to blebbistatin or Y27632 treatment (N=3, at least 15 cells per condition). Scale bars: 10 μm in overview and 2 μm in zoom-ins.

Paxillin phosphorylation has also been linked to vinculin recruitment (Pasapera et al., 2010). Our initial analysis indicated that vinculin is not organized in discrete clusters within focal adhesions (Fig. 1). However, this does not preclude that vinculin is relevant for the spatial organization of pPax. To test this, we cultured mouse embryonic fibroblasts from vinculin knockout mice (MEF vin -/-) and analyzed pPax-Y118 organization in these cells compared to MEF wt cells (Fig. 7). As shown before, paxillin appeared spread throughout focal adhesions while pPax-Y118 remained in separated clusters. Quantitative analysis of pPax-Y118 spot distance revealed an increase of cluster distance in absence of vinculin (Fig. 7E). Additionally, we tested the impact of cellular contractility in MEF cells (Fig. 7C, D) as we did before for REF cells (Fig. 6). These experiments again confirmed that reduced cellular contractility decreases pPax-Y118 spacing (Fig. 7E).

**Figure 7:**
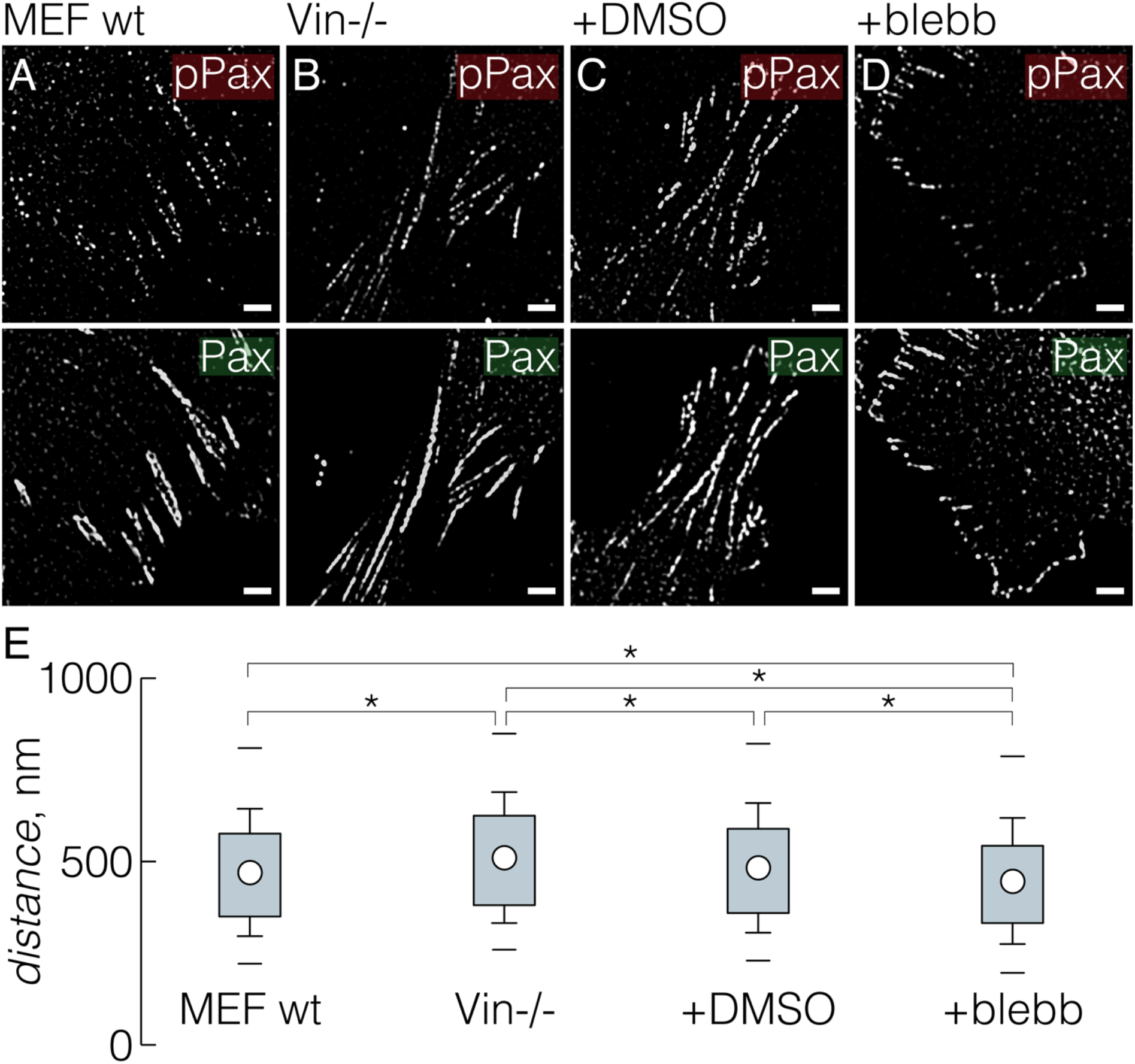
Cell contractility and vinculin expression modify the distance of pPax-Y118 clusters. (A) MEF wt cells were cultured on cover slips and stained for paxillin (green) and pPax-Y118 (red). (B) MEF cells from vinculin knockout mice (Vin -/-) were treated as described in (A). (C-D) Cells were treated as described in (A) but treated with DMSO or blebbistatin (20 μM) for the last hour of the experiment when indicated. (E) Quantification of pPax-Y118 cluster distance for cells described in (A-D) (N = 3, at least 20 cells were analyzed per condition). Scale bar: 2 μm.

To conclude, acto-myosin contractility affects the spacing of pPax-Y118 clusters within focal adhesions in REF cells and in MEF wt cells. Moreover, we could show that vinculin expression impacts pPax-Y118 cluster spacing. Vinculin is described to stabilize the connection between talin and actin fibers (Humphries et al., 2007). Thus, vinculin-dependent modulation of pPax-Y118 spacing again points to the relevance of actin-mediated cellular forces for the spatial organization of pPax.

## Discussion

We set out to study whether adhesome proteins show different lateral organization within integrin-mediated adhesions. We observed that several adhesome proteins are spread quite continuously throughout focal adhesions lacking any apparent organization. In stark contrast, FAK and proteins with phosphorylated tyrosines (shown with antibodies against pTyr, pPax-Y31, pPax-Y118, and pFAK) appeared organized in confined, discrete clusters within focal adhesions (see also Fig. 8). Super-resolution microscopy has been used before to study focal adhesions and substructures have been observed on different spatial levels. Hu and colleagues (Hu et al., 2015), as well as Young and Higgs (Young & Higgs, 2018) studied adhesions with similar resolution as we did. Both groups observed that classical focal adhesions, as observed with diffraction-limited microscopy, are in fact an assembly of parallel, but separated stripes (see also Fig. 8, zoom-in on the left). This organization can also be seen here in images of paxillin and talin-1 (Fig. 1). However, compared to the work by Hu et al. and Young & Higgs, we found an additional layer of spatial organization within adhesions: phosphorylated paxillin and FAK are restricted to discrete clusters within single stripes forming focal adhesions. We confirmed these pPax/FAK clusters with different antibodies, by recombinantly expressing FAK-GFP, in different cell lines, and with different microscopic techniques. This lateral focal adhesion structure remained so far, and to the best of our knowledge, undescribed. pPax/FAK clusters are 210 +/− 70 nm in diameter and are in average separated from each other by around 500 nm within single focal adhesion. Smaller lateral clusters within adhesions have also been described with pointillistic super-resolution techniques (Bachmann et al., 2016; Changede et al., 2015; Shroff et al., 2008; Shroff et al., 2007). However, these pointillistic clusters have been observed for integrins, vinculin or paxillin and were described to be below 100 nm in diameter. Thus, based on composition and structure size, it appears that pPax and FAK clusters described here are not identical to other clusters in focal adhesions mentioned so far.

**Figure 8:**
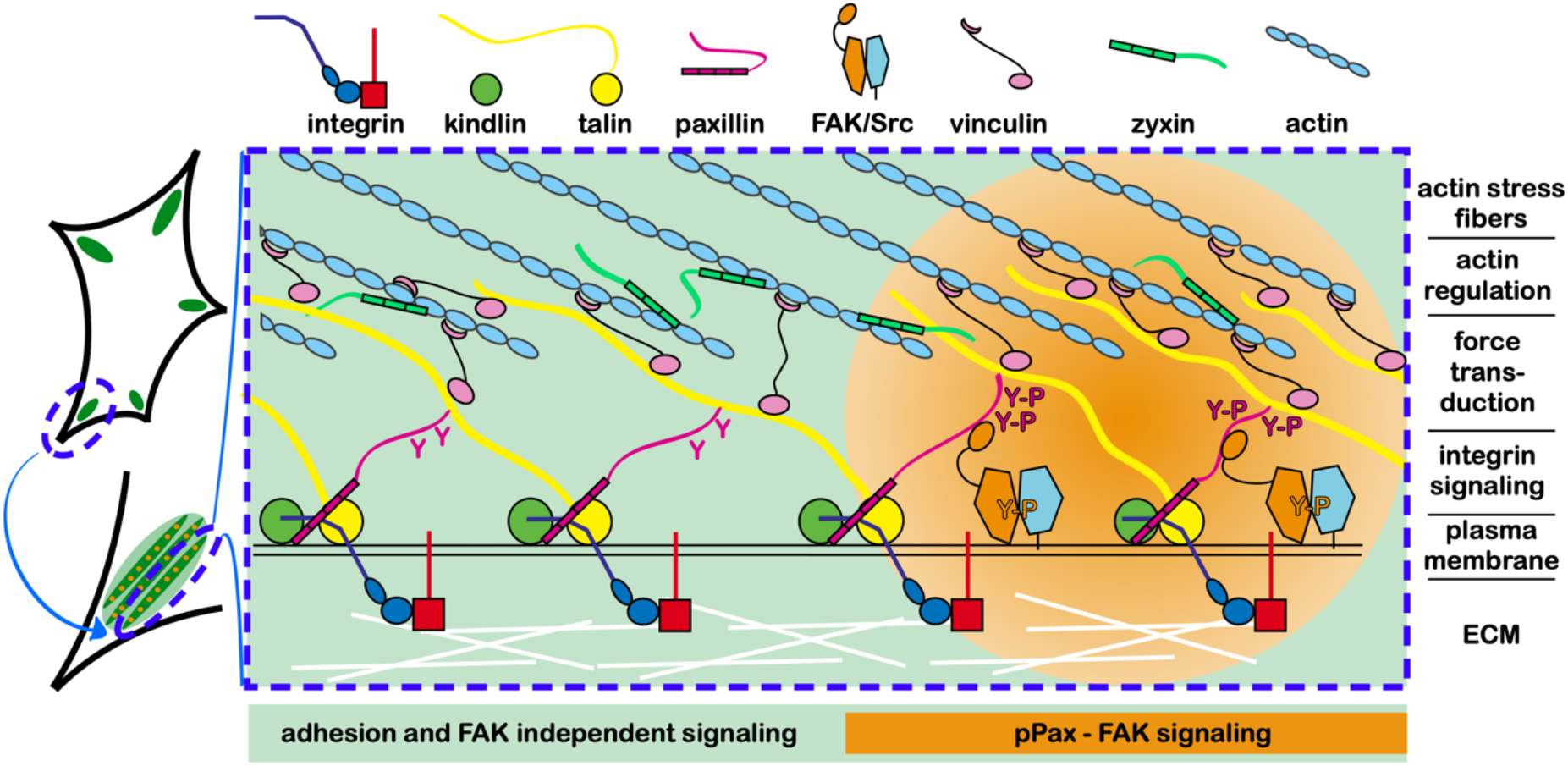
Focal adhesions have a lateral sub-structure of pPax/FAK spots while other adhesome components are found throughout adhesions. (Left) Increased microscopic resolution revealed that focal adhesions (green ellipse) consist of smaller, parallel strips (lower left: shown in dark green on green background; see also Fig. 1 and (Hu et al., 2015; Young & Higgs, 2018)). Additionally, we demonstrated here that pPax/FAK is organized in discrete spatial clusters within focal adhesions (orange spots in dark green focal adhesion stripes). (Central cartoon) These findings establish a lateral organization of focal adhesions with domains of pPax-FAK signaling (orange part in central cartoon; magenta ‘Y-P’ symbolizes pPax-Y31 and pPax-Y118, orange ‘Y-P’ symbolizes pFAK-Y397) while the remaining focal adhesion area (green) organizes FAK-independent signaling and adhesion. All other adhesome components we tested are laterally spread throughout adhesion without a restriction in spots as observed for pPax/FAK. At the same time, focal adhesions are organized in functional, axial layers (indicated on the right; see (Kanchanawong et al., 2010)).

What are functions of pPax/FAK clusters? pPax-Y31 and pPax-Y118 both recruit adapter proteins that eventually increase Rac1 and decrease RhoA signaling (Deakin & Turner, 2008) but also modify paxillin dynamics (Ripamonti et al., 2020). The impact on Rho-GTPase signaling contributes to more exploratory lamellipodia in the cell periphery. Paxillin phosphorylated at these sites was also discussed to recruit vinculin in a contractility-dependent manner (Pasapera et al., 2010). Indeed, we observed that chemically reducing cell contractility as well as a knockout of vinculin both affect pPax-Y118 spacing (Fig. 6, 7). Concerning the organization of pPax/FAK in separated clusters, it is interesting to consider alternative spatial organizations: Why is pPax/FAK not homogenously spread throughout focal adhesions like other adhesome proteins that we tested? One possibility might be that pPax/FAK signals have to be detected in a noisy environment of protein phosphorylation. In this regard, it might be advantageous to locally concentrate a pPax/FAK signal so that it may exceed background noise; this would not be the case if pPax/FAK was homogenously spread in focal adhesions. A mechanism of local enrichment to increase signal strength is so far hypothetical for focal adhesions. However, the general idea of local signal enhancement by concentrating the signal also applies to other cell signaling events. Lipid rafts concentrate transmembrane proteins, myelinated axons organize localized depolarization at myelin-sheath gaps, and filopodia concentrate signaling pathways in a limited volume compared to the complete cell body. Thus, it might not be surprising if signaling hubs like focal adhesions also implement the same strategy.

Another interesting question is the molecular mechanism that organizes and keeps pPax/FAK in clusters. Work by the Horwitz group indicated a preferred association of FAK with pPax-Y31 and pPax-Y118 (Choi et al., 2011; Digman et al., 2008). This could point to a positive feedback loop wherein FAK phosphorylates paxillin which in turn stabilizes FAK at this spatial position within a focal adhesion. This would also explain our observation that specifically pPax-Y31/118 and FAK/pFAK are organized in clusters. Such a local enrichment of FAK might leave other parts of the adhesion depleted of FAK (and of pPax-Y31 and pPax-Y118). This, in turn, would give rise to the periodic pattern of pPax/FAK spots within focal adhesions that we observed. Importantly, this mechanism is only dependent on paxillin recruitment to integrin-mediated adhesions and FAK activation. These are conditions that are easily fulfilled in adhesions: stable integrin-mediated adhesions cannot exist without paxillin and FAK is activated by many different pathways. A spot-creating mechanism like this, explaining pPax/FAK clusters without many prerequisites, would also explain why we observed pPax/FAK clusters very consistently in several cell lines and under different conditions. Concerning the impact of force-dependent changes in pPax spacing it is interesting to note that paxillin recruitment itself is contractility dependent (Bachmann et al., 2020) while FAK activation is also discussed to be force-sensitive (Tapial Martínez, López Navajas, & Lietha, 2020).

In summary, we described here new features of lateral adhesion organization based on super-resolution microscopy techniques. It was surprising to us how consistently pPax/(p)FAK/pTyr are organized in spatially separated clusters while some kind of “spotty appearance” of these stainings is surely well known to everyone working with these stainings. Importantly, however, we found this organization in clusters in different cell lines and overall to be conserved under a range of conditions. Moreover, experiments with reduced cellular contractility and modified vinculin expression indicate an active regulation of spot distance. Thus, we expect that this spatial organization will emerge as an important aspect of paxillin and FAK signaling of integrin-mediated adhesions.

## Acknowledgment

M. Bachmann acknowledges funding by Deutsche Forschungsgemeinschaft via Karlsruhe School of Optics and Photonics (KSOP) and BA 6471/1-1. A. Skripka acknowledges funding from the Erasmus Mundus Europhotonics program. The work of M. Bastmeyer is supported by the Deutsche Forschungsgemeinschaft (DFG, German Research Foundation) under Germany’s Excellence Strategy through EXC 2082/1-390761711 (Karlsruhe-Heidelberg 3DMM2O Excellence Cluster). The work of B. Wehrle-Haller is supported by the Swiss National Science Foundation (grant 310030_185261). We thank the bioimaging core facility of University of Geneva for their help and support.

## Material and Methods

### Cell culture

Cells were cultured in DMEM (Pan-Biotech, Germany) supplemented with 10% FCS (HyClone, USA) at 5% CO2 and 37 °C. NIH3T3 cells, HeLa cells and NRK cells were obtained from ATCC (USA). Rat embryonic fibroblasts were a gift from B. Geiger and human foreskin fibroblasts (HFF) were obtained from PromoCell (Germany). MEF wt cells and MEF from vinculin knockout mice (MEF Vin -/-) were a gift from W. H. Ziegler (Mierke et al., 2010). Transfections were carried out with Lipofectamine 2000 (ThermoFischer) according to manufacturer’s instructions. cDNA encoding full-length mouse β3-wt GFP integrin expressed in a cytomegalovirus promoter-driven pcDNA3/EGFP vector has been previously described (Ballestrem, Hinz, Imhof, & Wehrle-Haller, 2001), zyxin-RFP was a gift from A. Huttenlocher (addgene #26720, (Bhatt, Kaverina, Otey, & Huttenlocher, 2002)), and vinculin wt GFP was a gift from C. Ballestrem (Humphries et al., 2007).

### Antibodies and reagents

Inhibition experiments were performed with blebbistatin (Sigma-Aldrich, USA) or with the ROCK inhibitor Y27632 (Sigma-Aldrich, USA) at concentrations as indicated. Immunostaining was performed after fixation of cells with 4% PFA (Sigma-Aldrich, USA) in PBS. Reagents used for immunostaining were mouse antibodies against paxillin (BD Biosciences, #610051, 1:500 or as indicated), vinculin (Sigma-Aldrich, #V-9131, 1:100) or rat antibodies against β1-integrin (9EG7 clone, BD Transduction, #553715, 1:100) or rabbit antibodies against pPax-Y118 (Cell Signaling, #2541S, 1:500 or as indicated), for Supplementary Fig. 3 against pPax-Y118 (ThermoFischer, #44722G, as indicated) and against pPax-Y31 (ThermoFischer, #44720G, as indicated), against pTyr (Sigma-Aldrich, #T1325, 1:500), against pFAK-Y397 (ThermoFischer, #700255, 1:500). Primary antibody staining was followed by washing steps and incubation with antibodies against mouse labeled with Cy3 (Jackson Immunoresearch, #115-165-146, 1:500), against rabbit labeled with Alexa Fluor 488 (ThermoFischer, #A11070, 1:500) or Cy3 (1:500, Dianova, #111-165-144, 1:500), or with phalloidin coupled to Alexa Fluor 647 (1:200, ThermoFischer, #A22287). Primary rat antibodies were visualized with Alexa Fluor 488 labeled secondary antibodies (ThermoFischer, #A11006, 1:500).

### Microscopy

SIM imaging was performed on a non-serial Zeiss Elyra PS.1 microscope with a 63x/1.4NA oil immersion objective and an Andor iXon EMCCD camera. The grid for SIM was rotated three times and shifted five times leading to 15 frames raw data out of which a final SIM image was calculated with the structured illumination package of ZEN software (Zeiss). For SIM live cell microscopy, the incubation chamber was heated to 37°C. During imaging, cells were cultured in imaging medium (F12 + 25 mM HEPES + 200 mM L-glutamine + 1% penicillin/streptomycin + 10% FCS, pH 7.2). Values for calculation were selected for best resolution without causing image artifacts. Channels were aligned using a correction file that was generated by measuring channel misalignment of fluorescent tetraspecs (ThermoFischer, #T7280). TIRF pictures were taken with the same microscope and a 100x/1.46NA oil immersion objective. Confocal laser scanning imaging was performed on a Zeiss LSM 510 with a 63x/1.4NA oil immersion objective or with an Zeiss LSM 800 with the same objective and in super-resolution AiryScan mode.

### Image analysis

Images were prepared and analyzed (intensity profiles) with ImageJ software package (Schindelin, Rueden, Hiner, & Eliceiri, 2015). Line profiles were measured by averaging over 5 neighboring pixels (= 200 nm) in order to consider the full width of focal adhesion stripes. For analysis of spot-to-spot distance, we used a custom-written software, utilizing MATLAB Image Processing Toolbox (The MathWorks Inc., USA). Workflow: Two images, one serving as a focal adhesion mask and one being the staining of interest, are needed. Mask images were paxillin or vinculin stainings in our case (Fig. S1 – Mask) while pPax, pFAK, or pTyr were stainings of interest (Fig. S1 – Image of interest). Mask image was thresholded and structures below an area limit of 1-3 μm^2^ were excluded in order to limit the analysis to focal adhesions. Optionally, focal adhesions within a mask could be slightly broadened by adding desired number of pixels (typically around 1) around their periphery to ensure them covering most of stainings of interest within focal adhesions. Then, all focal adhesions within the mask were indexed and fitted as ellipses to extract their position, length, and width (Fig. S1 – characterize). Based on these features, stainings within corresponding focal adhesion were isolated, and an intensity line profile was created for a peak-to-peak distance calculation (Fig. S1 – Intensity profile). Size of individual spots was also extracted from intensity line profiles as width of each peak at its half prominence.

### Statistics

Data in box plots is represented as: dot within the box representing mean value, box outline representing 25^th^ and 75^th^ percentiles, whiskers representing standard deviation, and upper and lower bars representing 5^th^ and 95^th^ percentiles. Statistical comparisons are calculated with two-tailed Student’s t-test based on the number of independent experiments and between conditions as indicated in the figures (*: p < 0.05) Preparation of graphs and statistical significance testing were done with OriginPro 2017 software (OriginLab Corp., USA). All experiments were reproducible and were carried out as often as indicated in the figure legends. All pictures are representative examples from at least three independent experiments.

---

**Supplementary Fig. 1:**
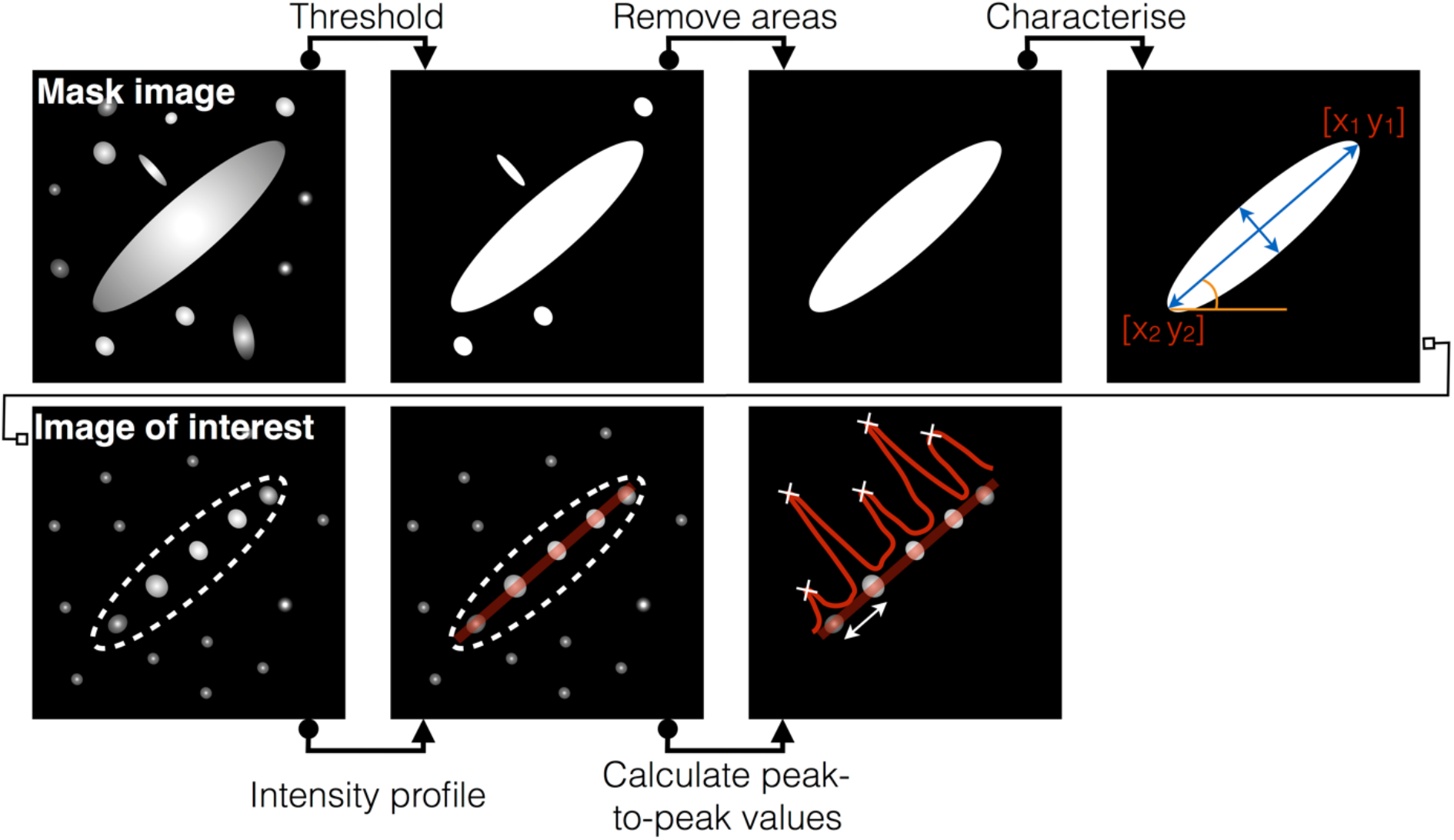
Scheme of the analysis for the spacing between pPax-Y118 clusters as shown in Figure 2. An image of paxillin staining (first row) is used as a mask for the subsequent analysis of the pPax-Y118 staining (second row). The paxillin staining is thresholded and objects smaller than focal adhesions (1-3 μm^2^) are discarded. This image is used as a mask for the corresponding pPax-Y118 staining (second row). Then, the intensity profile within the masked area is measured and the peaks of the intensity profile are calculated. The distance between these peaks yields the center-to-center distance of pPax-Y118 clusters.

**Supplementary Fig. 2:**
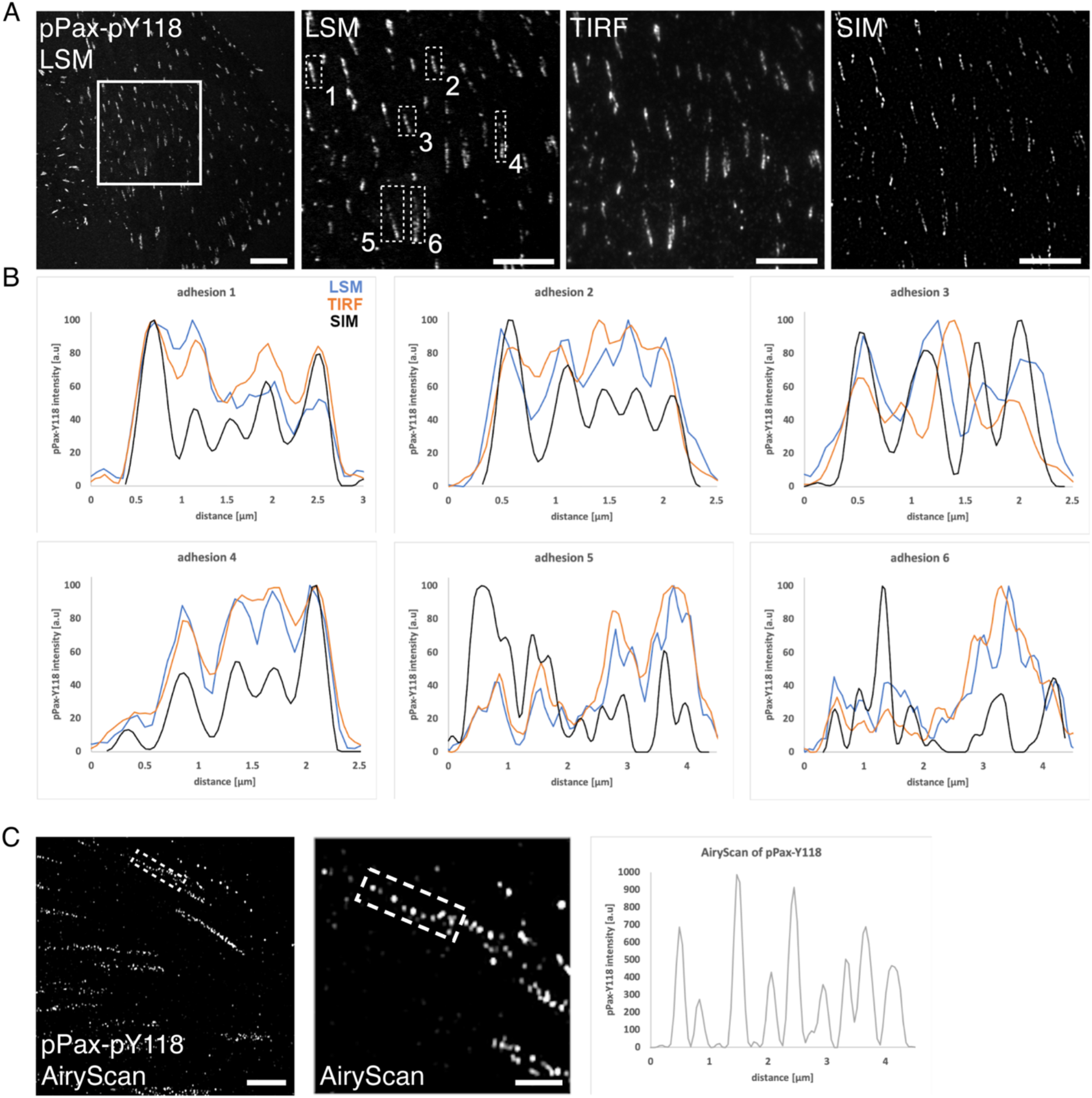
Super-resolution reveals a periodic pattern of paxillin phosphorylation. (A) REF cells were stained with indirect immunofluorescence for pPax-Y118. Samples were first imaged at a confocal laser scanning microscope (LSM) and then transferred to an Elyra PS.1 microscope for total internal reflection microscopy (TIRF) and SIM. The same cells already imaged with LSM were identified and imaged with TIRF and SIM. (B) Intensity profiles were measured for the same focal adhesions (dashed white boxes in zoom in for LSM in (A)) and plotted for every adhesion and the respective technique. Only SIM was reliably able to resolve the consistent organization of pPax in well separated clusters. However, some pPax-Y118 clusters could be resolved with either TIRF or LSM confirming that these spots are not artificial due to SIM technique. (C) Cell treated as described in (A) imaged with super-resolution mode of AiryScan on a ZEISS LSM 800. Intensity profile indicates separated pPax-Y118 clusters. Scale bars: 10 μm in overview in (A), 5 μm in zoom-ins in (A) and in overview in (C), 2 μm in zoom-in in (C).

**Supplementary Figure 3:**
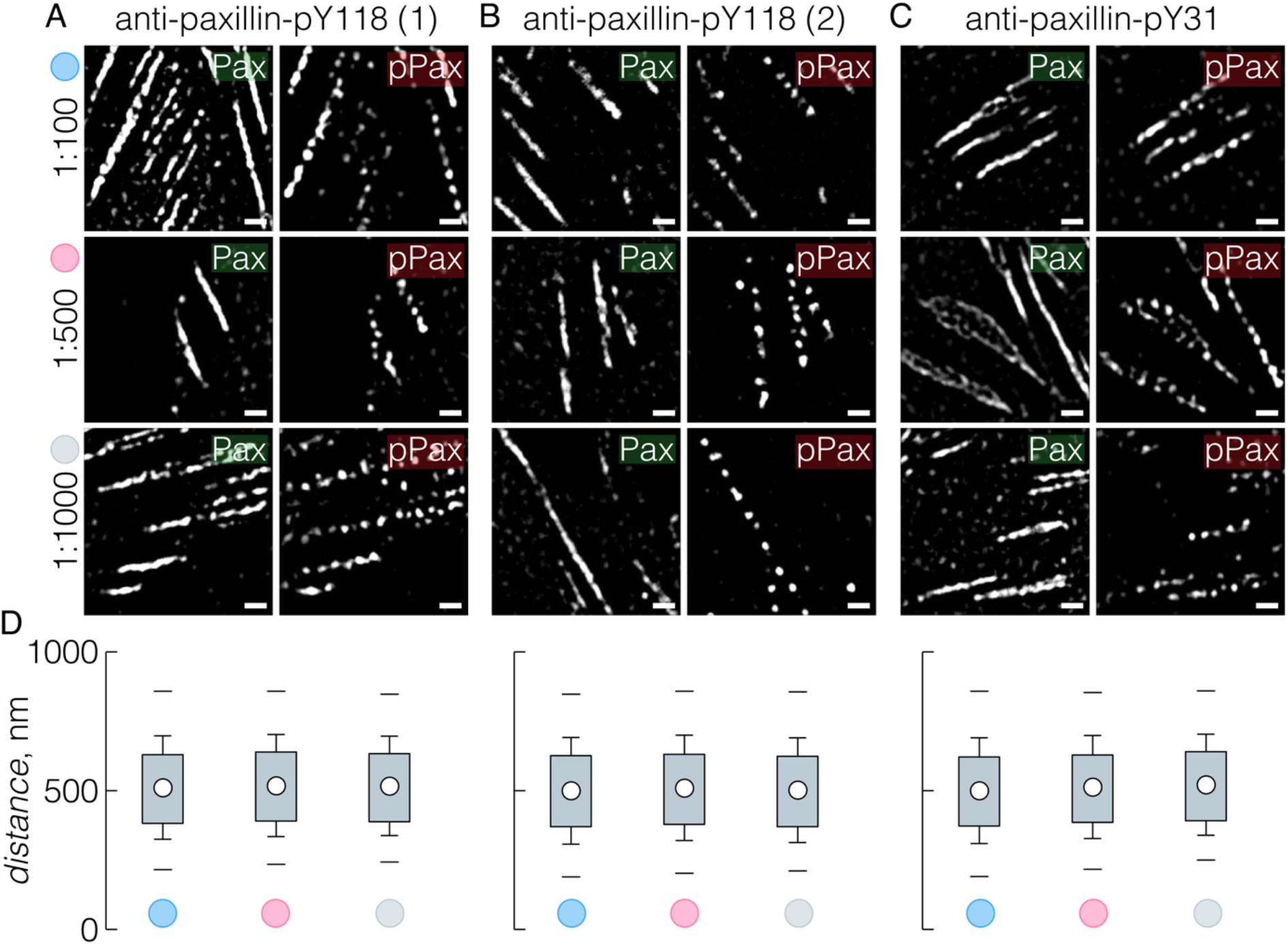
Organization of pPax-Y118 in spots is irrespective of primary antibodies and phosphorylation sites. REF cells were stained with primary antibodies for paxillin (green) with a dilution of 1:500 for all conditions. (A-B) pPax-Y118 was stained with two different antibodies; (A) antibody used throughout this study or (B) antibody from another distributor. (C) Staining for pPax-Y31. (A-C) Primary antibodies against pPax-Y118 or pPax-Y31 were diluted as indicated on the left. (D) Distance analysis for all different primary pPax antibodies and for all dilutions tested showed no significant effect of primary antibody or dilution (N = 3, n = 13). Scale bars: always 1 μm.

**Supplementary Figure 4:**
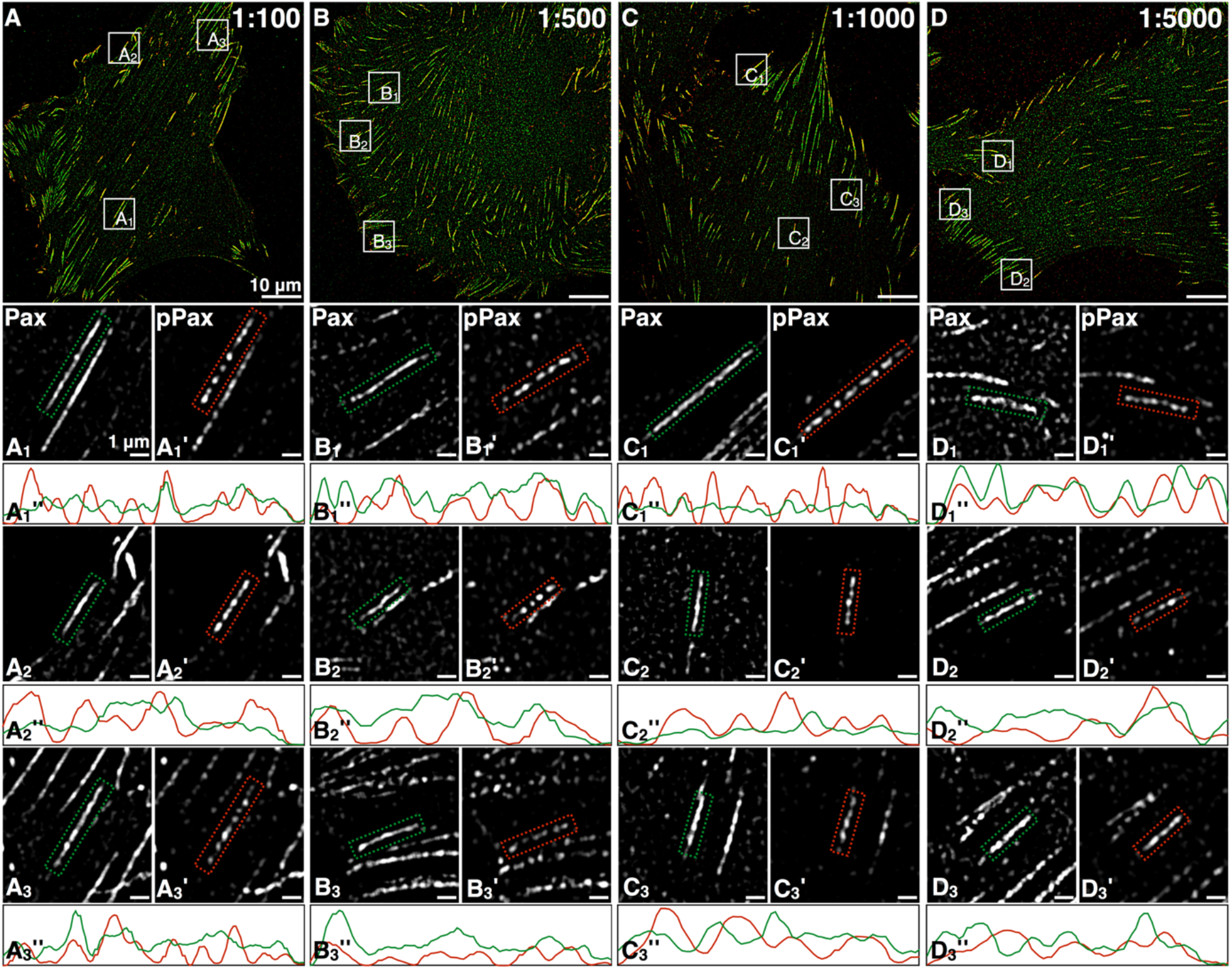
Paxillin shows no spot-like organization irrespective of antibody dilution. (A-D) We stained REF cells with primary antibodies for paxillin (green) and for pPax-Y118 (red). Primary antibodies for pPax-Y118 were always diluted 1:500. We varied the concentration for primary antibodies for paxillin to test for dilution dependent influences: (A) 1:100, (B) 1:500, (C) 1:1000, and (D) 1:5000. (A_1-3_-D_1-3_) Paxillin always appears homogeneous in focal adhesions while (A’_1-3_-D’_1-3_) pPax-Y118 appears in spots. (A_1-3_” – D_1-3_”) Intensity profiles for paxillin (green) and pPax (red) confirm this difference irrespective of the concentration of primary antibodies for paxillin. Scale bars: (A-D) 10 μm, (A_1-3_ – D_1-3_; A_1-3_’ – D_1-3_’) 1 μm.

**Supplementary Figure 5:**
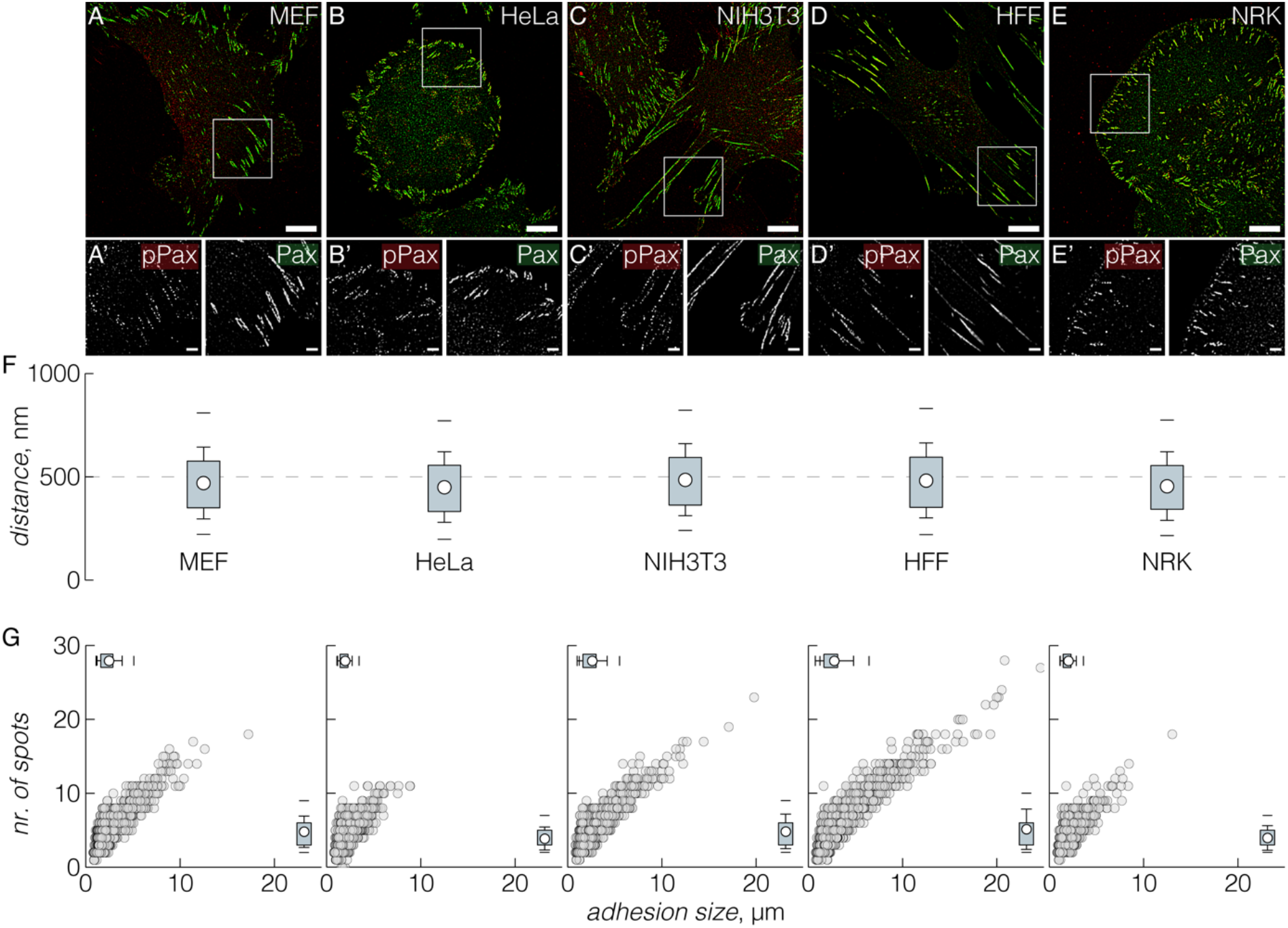
pPax-Y118 organizes in spots in several different cell lines. (A-E) Cancer cells (HeLa), mesenchymal cells (mouse fibroblasts NIH 3T3, mouse embryonic fibroblasts MEFs, and human foreskin fibroblasts HFF), and epithelial cells (normal rat kidney cells NRK) were stained with indirect immunostaining for paxillin (green) and pPax-Y118 (red). (A’-E’) Magnifications show a continuous organization of paxillin in focal adhesions while pPax-Y118-staining displayed an organization in distinct spots for every cell line tested. (F) Distance analysis of neighbored pPax-Y118 spots within focal adhesions and (G) spot number vs. adhesion length plot both revealed comparable organization of pPax spots within focal adhesions (N = 3, n = 18). Scale bars: 10 μm in all overview images and 2 μm in zoom-ins.

## List of abbreviations

SIM: super-resolution structured illumination microscopy
pPax: phosphorylated paxillin
pPax-Y31: phosphorylated paxillin at tyrosine 31
pPax-Y118: phosphorylated paxillin at tyrosine 118
FAK: focal adhesion kinase
pFAK: FAK phosphorylated at tyrosine 397
pTyr: phosphorylated tyrosine
REF: rat embryonic fibroblast
HFF: human foreskin fibroblast
HeLa: cancer cell line
NRK: epithelial cell line
NIH 3T3: fibroblastoid cells
MEF: mouse embryonic fibroblasts
MEF vin -/-: vinculin knockout (-/-) cells

